# Distinct role of flexible and stable encoding in sequential working memory

**DOI:** 10.1101/525220

**Authors:** Hyeonsu Lee, Woochul Choi, Youngjin Park, Se-Bum Paik

## Abstract

The serial-position effect in working memory is considered important for studying how a sequence of sensory information can be retained and manipulated simultaneously in neural memory circuits. Here, via a precise analysis of the primacy and recency effects in human psychophysical experiments, we propose that stable and flexible coding take distinct roles of retaining and updating information in working memory, and that their combination induces serial-position effects spontaneously. We found that stable encoding retains memory to induce the primacy effect, while flexible encoding used for learning new inputs induces the recency effect. A model simulation based on human data, confirmed that a neural network with both flexible and stable synapses could reproduce the major characteristics of serial-position effects. Our new prediction, that the control of resource allocation by flexible-stable coding balance can modulate memory performance in sequence-specific manner, was supported by pre-cued memory performance data in humans.

## Introduction

The brain receives various types of sensory information from the external environment and encodes them as a form of working memory^1–4^. This enables short-term storage of received information and manipulation of it at the same time, which is crucial to cognitive processes such as visual and auditory perception of sequential information^5–7^.

Early studies reported that the capacity of working memory is limited^3,7,8^. Conceptual models suggested that working memory has a fixed number of slots, such as Miller’s magical number seven^9^ or Cowan’s number four^10^. More recently, psychophysical observations of working memory in multi-item tasks revealed that human working memory can be better described by the resource model where a limited memory resource is flexibly allocated to the information of each item so that the amount of allocated resource determines the memory resolution^3,11–14^. However, these conceptual models simply describe a relationship between memory performance and resource allocation, but do not account for the underlying principle of memory resource allocation that enables retaining and updating information in working memory.

One important observation in the sequential working memory task is that performance for each item varies by the order of presentation, referred to as the serial-position effect^5,15–20^ The performance curves of subjects typically appear U-shaped in consequence, because most subjects better memorize items presented first and last in the sequence than the others in the middle. These are often referred to as the primacy^17,18,21^ and recency effects^5,17,18,21^, respectively, and are considered to reflect a key mechanism of how neural resource is utilized, specifically in sequential memory coding.

Various models have been proposed to explain the underlying mechanism of this serial-position effect^18–20^, but a complete accounting of the observed results has not yet been achieved. For instance, one model suggested that the serial-position effects arise from the processes of temporal decay and restoration of memory^22,23^, but other studies claimed that a variation of retention time alone could not regenerate the observed profile of memory performance^5,24^. Similarly, another model suggested that the recency effect is explained by assuming a specific type of resource reallocation to recent items^3,5^, but the primacy effect could not be addressed together in this model. It also has been suggested that the primacy and recency effects could arise from declining encoding strength accompanied by response suppression during memory recall^24,25^, but the neural mechanism of this conceptual memory processes is not yet fully understood.

Here, we propose that the serial-position effect arises from two distinct types of neural encoding that are indispensable for working memory function. Stable encoding of information allows retaining of previous memory and results in the primacy effect; while flexible encoding enables update of recent memory and results in the recency effect. Our results not only explain the origin of the serial-position effects, but also suggest that coexistence of flexible and stable coding is required to form working memory.

First, we performed a human psychophysical experiment and precisely investigated the serial-position effect. Based on the quantitative analysis of order-dependent memory performance, we suspected that the primacy and the recency effect could arise from two different mechanisms. With a model neural network simulation of controlled synaptic plasticity, we found that stable synapses could retain old information, while flexible synapses could encode new information. Taken together, we could reproduce the observed serial-position effect by balancing the contribution of flexible and stable synapses in a model neural network. Our model also predicted that modulation of the flexible/stable synapse ratio would change the strength of the recency/primacy effects, and also modulate memory performance in an order-specific manner. Our prediction was validated by human psychophysical experiments, in which a pre-cue of stimulus information altered subjects’ performance order-specifically as predicted.

In summary, we propose that the serial-position effect of human sequential memory reflects distinct roles of flexible and stable neural encodings, and that this enables storage and instantaneous manipulation of information in working memory.

## Results

### Serial-position effects of sequential working memory

To quantify the serial-position effects of sequential working memory, we designed a human psychophysical experiment using non-semantic visual patterns of smoothed white noises to minimize any correlation between items (see Methods for details). Subjects were asked to memorize visual patterns presented sequentially and to recall the memorized sequence freely (Fig. 1a and Supplementary Fig. S1). As expected, a strong serial-position effect was observed in most subjects, in which memory performance for the first and last items in a sequence was higher than that for the other items (Fig. 1 b and c).

**Figure 1.**
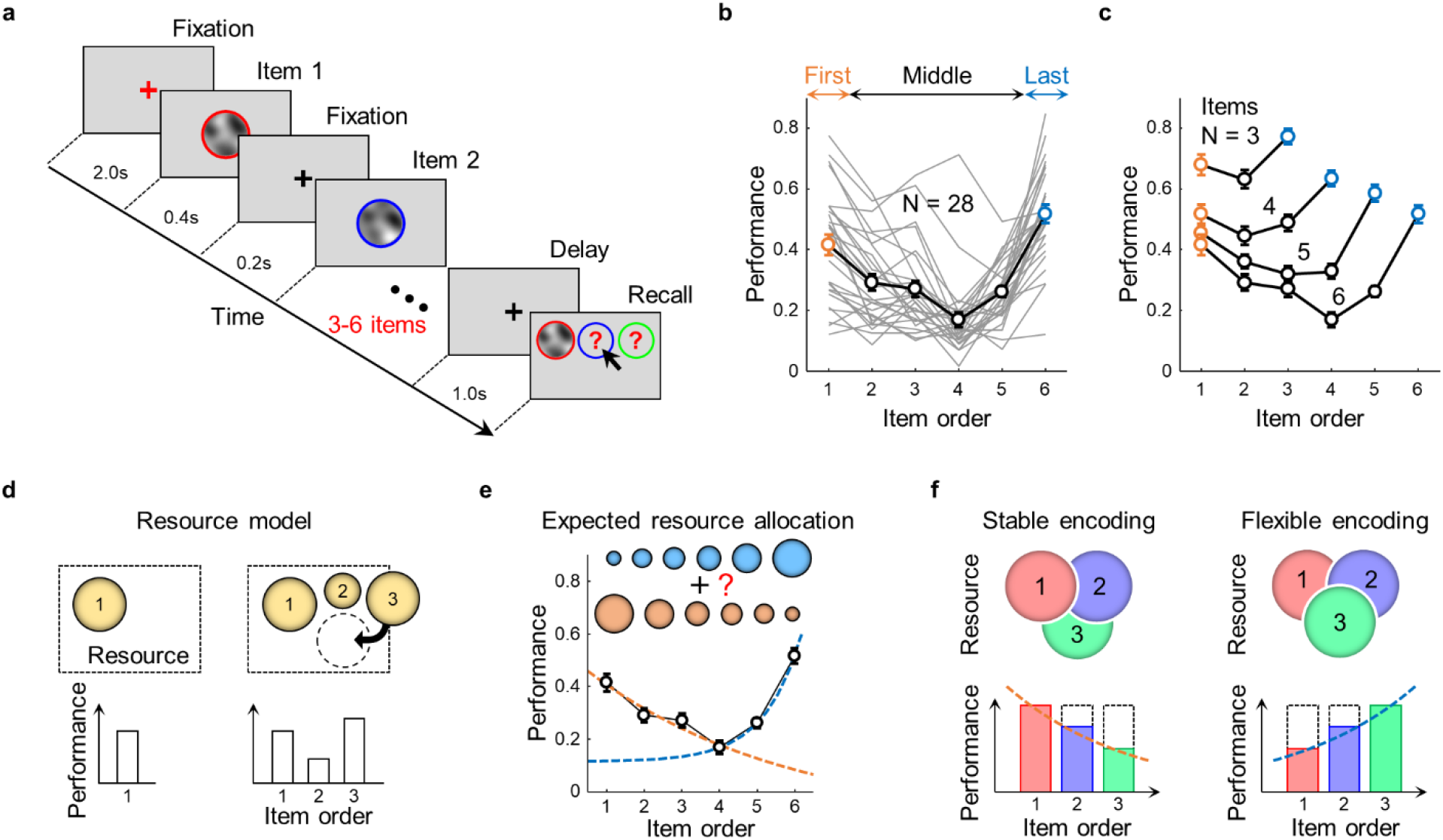
Recency and primacy effect of sequential working memory task. **a**, Experimental design for a sequential memory task. Subjects were asked to memorize visual patterns (N_items_ = 3-6) presented sequentially and to recall freely. **b**, Sample memory performance curve by item order (N_items_ = 6). Performance for the first (orange) and last (blue) items was higher than for the others (mean ± s.e.m.). Gray lines represent individual performance curves. **c**, Average memory performance curves. The serial-position effect was observed regardless of the number of items in a sequence (mean±s.e.m.; N_items_ = 3–6). **d**, Illustration of the resource model. The model assumes that the amount of allocated resources determines the memory performance for each item. e, Two types of resource allocation. The serial-position effect can be described with a decreasing (orange) and increasing (blue) resource allocation model. The primacy and recency effects were fitted with exponential functions, respectively (*y* = *aexp*(*bx*) + *c; a* = 0.64, *b* = −0.19,*c* = −0.12 for primacy effect, *a* = 9.37 × 10^-4^,*b* = 1.01,*c* = 0.12 for recency effect). **f**, Model hypotheses of stable and flexible encoding. We hypothesized that stable encoding would induce the primacy effect and that flexible encoding would induce the recency effect.

According to the resource model^3,5,26^, the amount of memory resource allocated to each item determines the performance (Fig. 1d). In this view, more resources need to be allocated to the first and last items to reproduce the serial-position effect in our observation. However, the U-shaped memory performance cannot be regenerated by assuming a simple form of resource allocation that monotonically increases or decreases. Instead, we hypothesized that the primacy and recency effects might arise from two distinct mechanisms of resource allocation (Fig. 1e): one with increasing amount of resources by order, and the other with decreasing. We observed that the primacy effect was well fitted to an exponential function decreasing by order, while the recency effect was to an increasing one. This suggests that two distinct types of resource allocation model are required to reproduce a complete profile of the serial-position effect.

In the primacy effect, decreasing performance suggests that the amount of resources allocated to each item decreases by order (Fig. 1f, left). This phenomenon can be explained if we introduce a scenario of “stable” coding of information, in which the resource used by an old item is very stable, so that it cannot be shared by a new item received. Then, an old item is better retained than a new one, the amount of resource decreases by order in this instance. On the other hand, increasing performance in the recency effect can arise from a “flexible” coding of information, in which the resource allocated to an old item can be readily overwritten by the information of a new item (Fig. 1f, right). Thus, old items are better retained than a new one in the stable coding scenario, while the memory of a previous item is degraded when a new item is memorized in the flexible coding scenario. Under these assumptions, we supposed that stable encoding would induce the primacy effect, while flexible encoding would induce the recency effect, and that the serial-position effect reflects a collaboration of the flexible and stable encodings in working memory. Thus, we modeled how memory resource is allocated under flexible and stable encoding schemes, by quantitatively analyzing the recency effect and primacy effect, respectively.

### Recency effect by flexible encoding

To model the profile of resource allocation by flexible encoding, we first examined how the performance for previous items was altered when a new item was introduced (Fig. 2a). For instance, we investigated how performance (or presumably the amount of allocated resource) for the previous three items is modulated by a new (fourth) item, by measuring difference between two performance curves (ΔPerformance) of which the number of total item is N = 4 vs. N = 3 (Fig. 2b). We found that performance for previous items was decreased by a new item, in a way that the correct ratio for more recent items was decreased more. Interestingly, the trend of memory degradation was observed to be similar in different cases (N = 6 vs. N = 5, N = 5 vs. N = 4 and N = 4 vs. N = 3) (Fig. 2c). This common trend of the performance change, normalized to the performance of the last item, was well fitted to a single power-law function (*y* = -*y*^|*x*|^; *γ* = 0.55), suggesting that the memory resource for previous items was taken by a new item with a constant ratio (*γ*).

**Figure 2.**
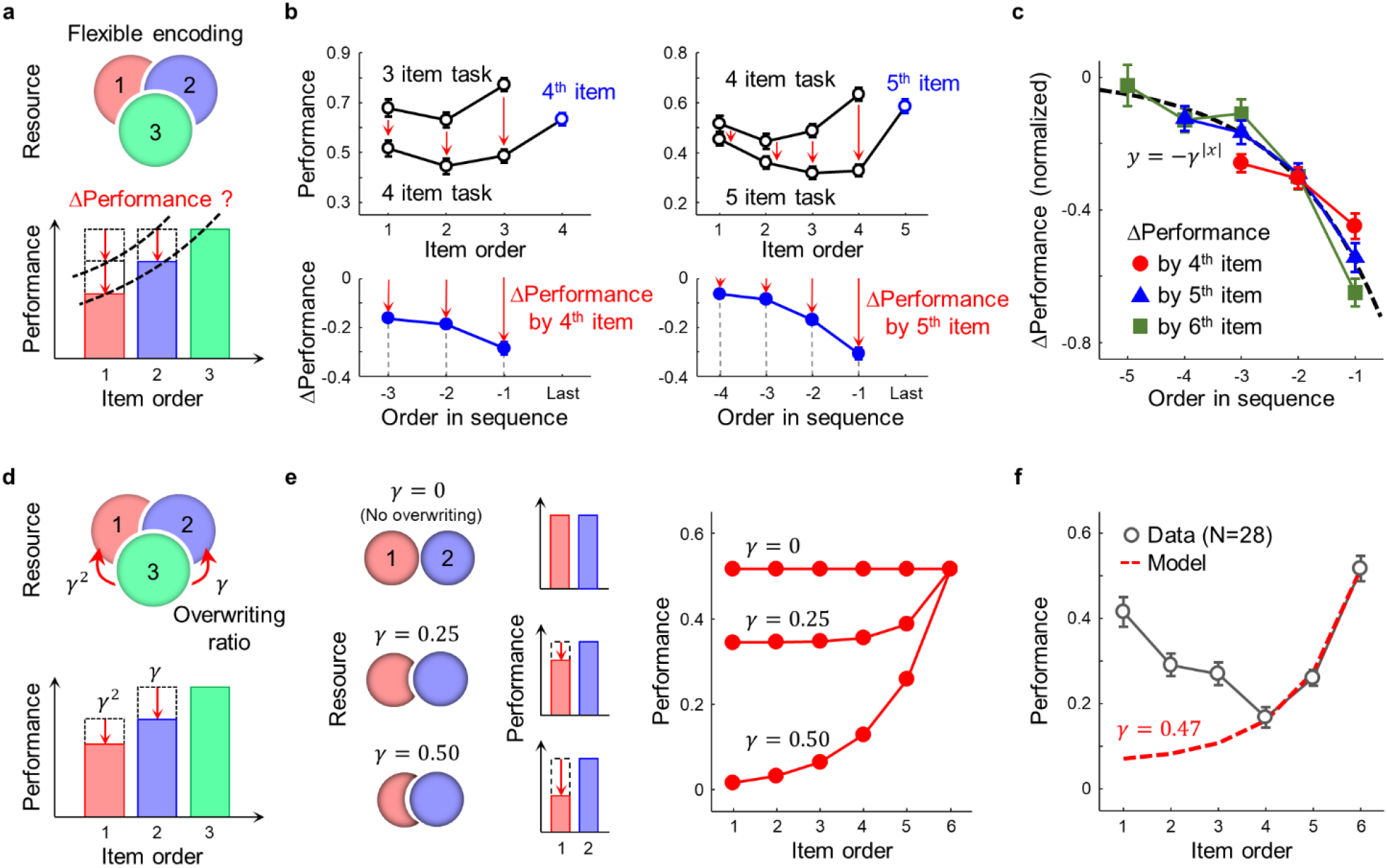
Flexible encoding induces the recency effect with resource overwriting. **a**, Illustration of flexible encoding. Under flexible encoding scheme, performance for old items (red and blue) is degraded by a new item (green). **b**, Estimation of resource overwriting by flexible encoding in data. Performance for previous items decreased when the last item was given (top; mean ± s.e.m.), and Δ Performance was better in recent items (bottom). c, Universality in performance change. Performance differences (normalized to performance for the last item) were well fitted with a single exponential function (*R*^2^ = 0.91, *y* = -*γ*^|*x*|^, *γ* = 0.55). **d**, Concept of sequential overwriting model. When a new item (green) is given, a new item overwrites memory resources of old items (red and blue) by a constant overwriting ratio (*γ*). Overwriting was assumed to follow a power-law function, following the observation in **c. e**, Performance curve simulated by a model. The recency effect, higher performance of recent items than for older items, was strengthened as the overwriting ratio increased. **f**, A fitted recency effect curve with sequential overwriting model. Memory performance for the last three items was fitted by error minimization (*γ* = 0.47). All error bars represent s.e.m.

Based on this observation that a new item overwrites the memory resource of older items, we proposed a sequential overwriting model as a revision of the resource model previously suggested^3,5,26^ (Fig. 2d). First, following the resource model, we assumed that an item is memorized in the memory resource allocated to each item and that the memory performance for each item is proportional to the amount of resource allocated. Next, we hypothesized that memory resource (or performance) for previous items are degraded by a new item with an overwriting ratio, *γ*(Fig. 2c and d), so that the memory resource decreases as a power of *γ* by the item order, as observed in our experiment. In this model, the amount of performance change in previous items by sequential overwriting can be estimated mathematically. After sequential overwriting of every item, memory performance was estimated by the amount of resource remaining (see Methods for details). In this scenario, the profile of the memory performance curve only varied by overwriting ratio, *γ*(Fig. 2e, left). For non-zero *γ*, memory performance for a recent item was always better than for previous items, signifying the recency effect. In addition, performance for the last item was not affected by resource overwriting (Fig. 2e, right). We found that our sequential overwriting model could not only reproduce the profile of the observed recency effect, but could also predict the degree of resource overwriting (estimated parameter *γ* = 0.47) that matches the observed profile of performance curve (Fig. 2f). Overall, our model implies that the recency effect might be a result of flexible encoding of sequential information.

### Primacy effect by stable encoding

Flexible encoding model alone cannot explain the other face of sequential working memory. The primacy effect reveals that allocated memory resource seems to decrease by order (Fig. 3a). To model the mechanism of declining memory resources, we introduced the concept of stable encoding of information, in which the resource allocated to an old item is very stable, so that it is not affected by a new item received (Fig. 3b). In this scenario, the amount of allocated resource decreases by order, because the total resource available is limited. Thus old items are better retained than a new one. To investigate this issue in the data, we questioned how the amount of allocated memory resource is determined when there is no influence of resource overwriting. For this, we investigated memory performance for the last items in various set sizes, because they are not affected by the next item in the sequence, even under flexible encoding scenarios (Fig. 3c). We observed that memory performance for the last item in an experiment decreases as set size increases (Fig. 3d). Thus, we inferred that the profile of memory performance unaffected by resource overwriting is a monotonously decreasing curve and that more resource is allocated for earlier items if there is no resource overwriting, consistent with our stable encoding scenario. From this profile of memory performance for the last items, we could estimate the relative amount of resource allocated to each item, which well fit an exponential function (*y* = *a* + *be*^-*c*(*x-d*)^, *a* = 0.45, *b* = 1.30, *c* = 0.50, *d* = 0.16).

**Figure 3.**
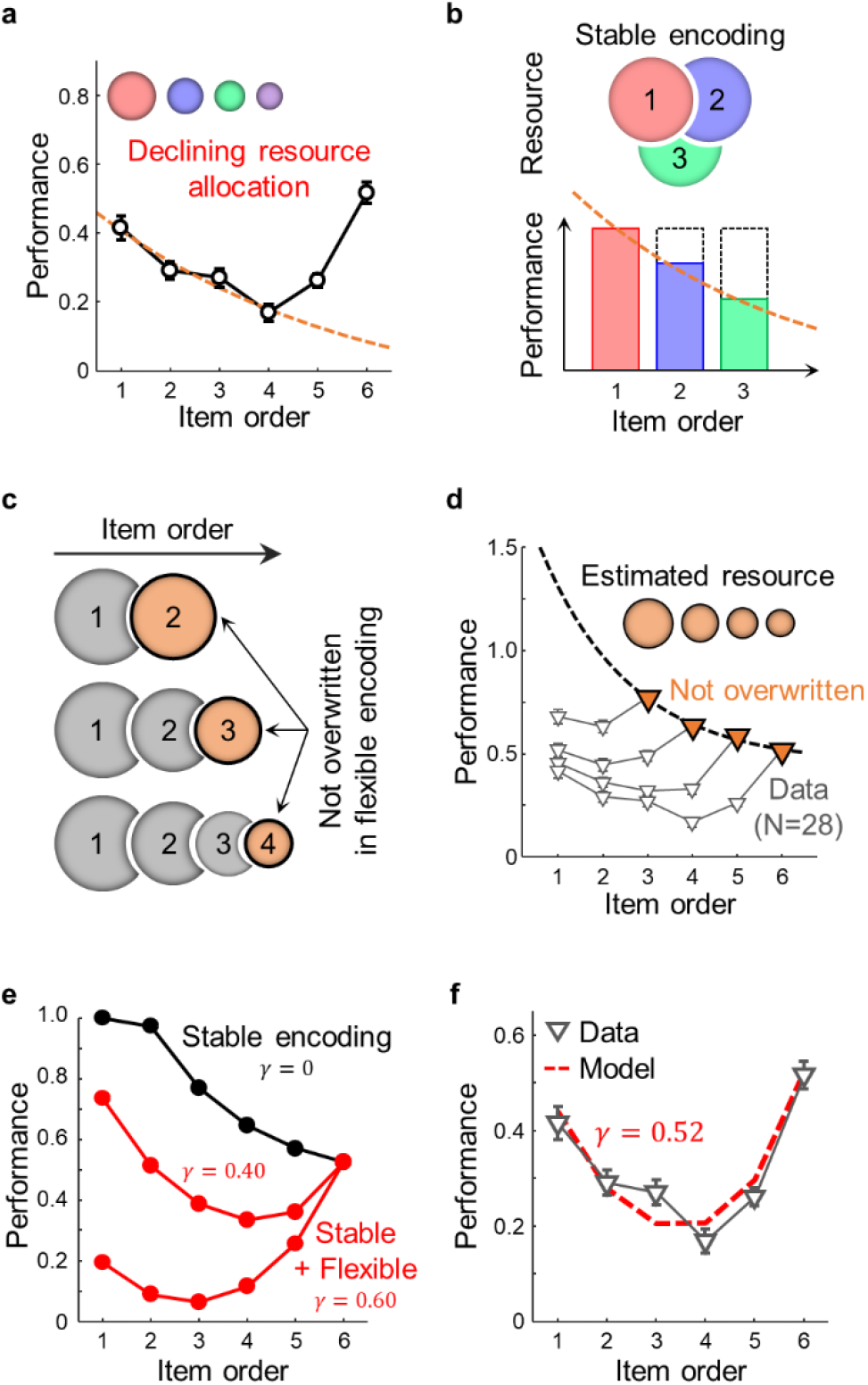
Stable encoding induces the primacy effect and coexistence of stable and flexible encoding generates the observed serial-position effect. **a**, Prediction for primacy effect. To induce the primacy effect, the amount of allocated resources needs to decrease by order (Inset). More resources are allocated to early-presented items. **b**, Illustration of stable encoding. Old items (red and blue) are better retained than a new item (green), because more resource is occupied by old items. **c**, Estimation of resources not affected by resource overwriting. Even in flexible encoding scheme, the last item is not overwritten by the other items (orange). d, Estimation of resource amount from data. The amount of resource allocated with no overwriting effect was estimated from memory performance of the last items (orange triangles), fitted to an exponential function (*R*^2^ = 0.99, *y* = *a* + *b exp*(-*c*(*x* – *d*)), *a* = 0.45, *b* = 1.30, *c* = 0.50, *d* = 0.16). **e**, Performance curves modeled with both stable and flexible encodings. Both primacy and recency effect were generated if both stable and flexible encoding contributed (red). **f**, Serial-position effect in sequential overwriting model fitted to data. The degree of overwriting was estimated by minimizing the mean squared error between the performance curves of model and data (*γ* = 0.52; mean±s.e.m.).

Based on these two scenarios of increasing and declining resource profiles, we hypothesized that working memory has both stable and flexible types of coding scheme. To model this idea, we performed a simulation to achieve a memory performance curve to which stable and flexible resources contributed together. We started from the observed profile of declining memory resources by stable encoding in Fig. 3d (only the primacy effect observed) and then added the flexible encoding component by allowing resource overwriting (*γ*>0), as modeled in Fig. 2e. We confirmed that both the primacy and recency effects can be observed only when flexible encoding (non-zero resource overwriting) was added to stable encoding (Fig. 3e). To reproduce quantitatively the serial-position effects observed in the experimental data, we performed a parameter search for the overwriting ratio by minimizing the error between the performance curves of model and data (Fig. 3f; see Methods for details). The model performance curve fitted (*γ* = 0.52) to the experimental observations suggested that the observed serial-position effect can arise when approximately half the memory resource of a new item affects previous items. Taken together, our model suggests that both flexible and stable encoding are required to generate the observed serial-position effect, both of which are also required, in principle, for working memory.

### Working memory simulation with flexible-stable model synapses

We further assumed that the memory performance curve for sequential information could be altered by the balance between the relative amount of stable and flexible resources in neural memory circuits (Fig. 4a). Our model predicted that if the whole resource performs flexible encoding only, a new item always overwrites old items. Thus, there would be the recency effect only (Fig. 4a, left). In the same way, if the whole resource was of the stable type, old items would always be better retained than a new item and strong primacy effect would be observed (Fig. 4a, middle). Therefore, to induce the serial-position effect, both sequential overwriting and declining resources must contribute together through the performance of flexible and stable encoding, respectively (Fig. 4a, right).

**Figure 4.**
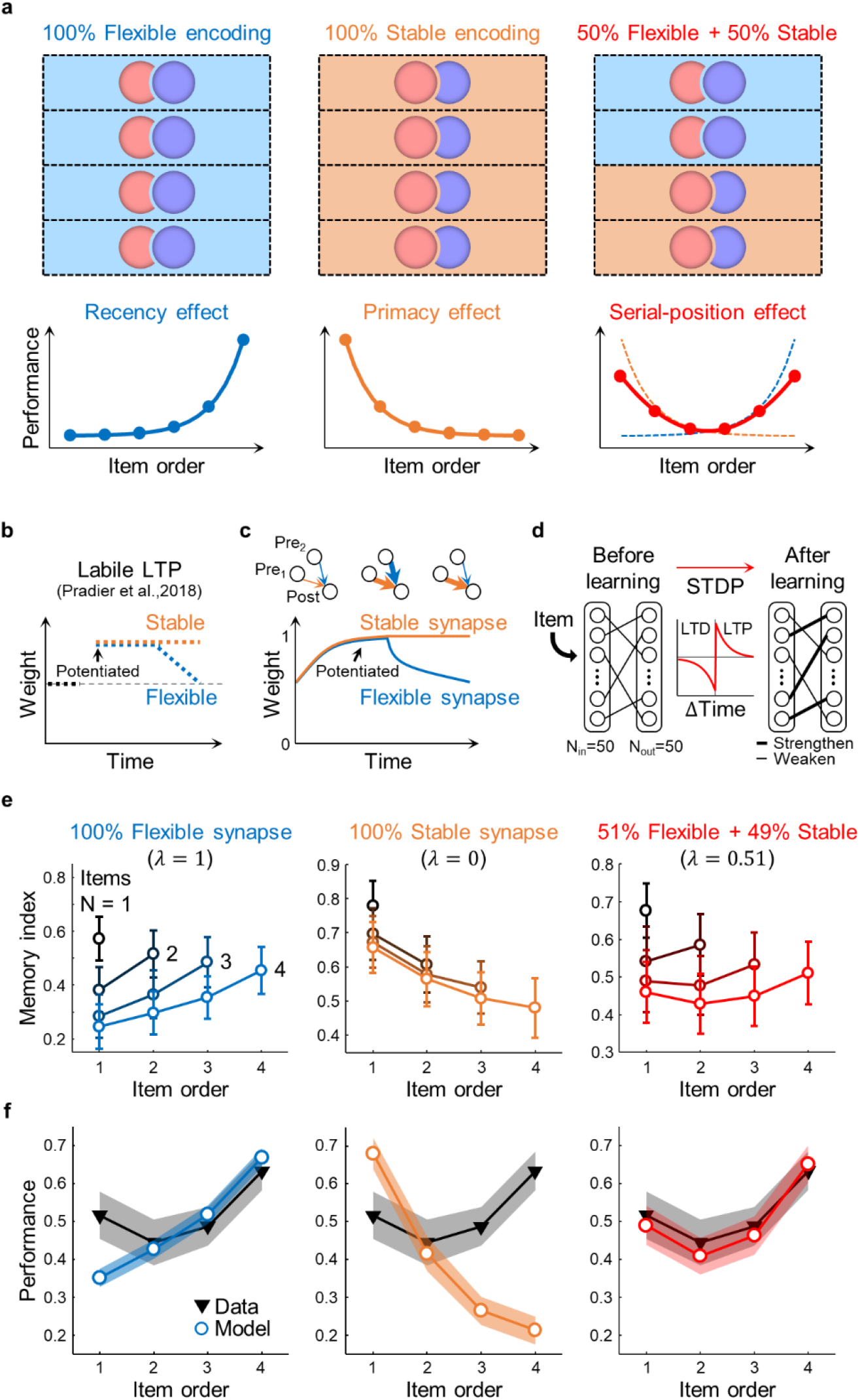
Flexible and stable synapses in model network induce flexible and stable encoding. **a**, Population model of stable and flexible encoding. (top) Each dashed box represents sub-regions of memory space where memory resource can be allocated. When an item is encoded, the memory resource for that item is allocated in memory space either flexibly (blue) or stably (orange). (bottom) Predicted memory performance of models in each condition. **b**, Illustration of the input frequency-dependent synaptic plasticity (labile LTP^38^). Potentiation remains stable (orange) or is reset rapidly (blue). **c**, Design of stable and flexible synapses. Synaptic weight of connection can be either increased (LTP) or decreased (LTD) during learning. For stable synapse (orange), its weight maintains stable after the weight is saturated. For flexible synapse (blue), its weight continuously changes during learning. **d**, Design of a feedforward neural network for memory simulation. The network consists of two-layers: input layer and output layer (50 neurons, each). Synaptic weights of connections can be updated using the STDP learning rule. **e**, Memory index change for sequentially given inputs. Four items (temporal spike patterns) were sequentially encoded to the network. The networks consist of flexible synapse only (left), stable synapse only (middle), or both flexible and stable synapses (right). **f**, Simulated memory performance. The recency and primacy effects were observed under the flexible and stable synapse cases, respectively (left, middle). A complete serial-position effect is observed only when both flexible and stable synapses coexist (right). Shaded area represents 95% confidence intervals.

So far, we have shown that collaboration of flexible and stable encoding can generate the serial-position effect, using a conceptual model only. If so, then what kind of neural factors can implement two distinct encoding schemes in a neural circuit? Previously, the conventional and predominant view has been that sustained activity of neurons during the delay periods is the neuronal basis of working memory representation^4,27–31^. However, more recent studies have suggested a dynamic coding scenario, in which short-term retention of information is patterned in neural activities via synaptic plasticity^32–36^. It was reported that neurons in rat prefrontal cortex (PFC) exhibit large heterogeneity in their intrinsic temporal stability so that some neurons retain stimulus information while others code more transient selectivity functions. This enables reconciling of persistent and dynamic coding of the working memory^37^. Similarly, we assumed that information processing achieved by the combination of stable-flexible encoding might be a key mechanism for understanding the neural basis of sequential working memory.

To propose a possible neural basis of the serial-position effect in working memory, we studied to determine if our conceptual model of stable-flexible encoding could be realized in a model neural circuit, by simply introducing stable/flexible components of synaptic plasticity. For this, we adapted a particular form of synaptic plasticity recently found, the labile longterm potentiation (LTP) (Fig. 4b)^38^, which can switch between stable and flexible encoding depending on conditions. This synapse potentiated by high frequency stimulation can be either maintained (stable) or depotentiated (flexible), depending on background activity frequencies. By adapting the dynamics of this labile LTP, we introduced two types of synapses into the model network: flexible and stable ones (Fig. 4c). The strength (weight) of the flexible synapse is allowed to continuously change during learning, so that the synapses can learn new information by sacrificing old information. In contrast, a stable synapse was set not to change its synaptic weight, once it was potentiated or depotentiated enough to a certain threshold value (see Methods for details).

Under this model condition, we expected that flexible and stable synapses could induce the recency and primacy effects, respectively, and that a mixed population of them could reproduce the observed serial-position effect. To test this idea, we made a model neural network that received random spike trains as input, and for which the feedforward wirings between input and output layers could be trained using the spike-timing-dependent plasticity (STDP) learning rule (Fig. 4d and Supplementary Fig. S2a)^39^. Performance (Memory index) of the trained network was defined as the consistency of response activity of the network to each trained input pattern, and was measured by the average pairwise cross-correlations between the binary output firing patterns of response activity, similar to those in hippocampal engram studies of fear memory^40,41^ (see previous Methods^39^ for details, and Supplementary Fig. S2b). Thus, with ‘1’ as the memory index, if the neural output pattern for a particular input pattern was always the same for that input, it would mean that this pattern was completely memorized. On the other hand, if the memory index were ‘0’, it would mean that the network did not memorize an input pattern so that it generated a random response pattern. Using this simplified model, we compared the memory performance of the neural populations under three conditions at different rates of flexible synapses, *λ*: when the neural wirings consist of (1) flexible synapses only (*λ* = 1), (2) stable synapses only (*λ* = 0), and (3) both flexible and stable synapses (0 < *λ* < 1) (Fig. 4e and f).

When the network consisted of flexible synapses only, the memory index of newer items was higher than that of previous ones, regardless of the length of sequence (Fig. 4e, left). As in our previous conceptual model, flexible synaptic connections that encoded the information of the old items could be altered by information about new items in this case (sequential overwriting); thus this model condition generated the recency effect of working memory (Fig. 4f, left). In contrast, when the network consisted of stable synapses only, the memory index of newer items was lower than the old ones (Fig. 4e, middle). Stable synaptic connections that encoded the information of old items were unchanged during the learning of new items, thus later items had smaller numbers of synapses available for encoding (declining resources), leading to the primacy effect (Fig. 4f, middle).

When the network consisted of both types of synapses, characteristics of flexible and stable encoding were observed simultaneously (Fig. 4e, right). Old items were better retained than newer ones (primacy effect) early in the sequence, while newer items were better memorized than old ones (recency effect) later in the sequence. To match quantitatively the profile of human data, we performed a parameter search for a ratio between flexible and stable synapses in the network that would minimize the error between the model and data performance curves (*λ* = 0.51). As a result, the model could successfully generate both the primacy and recency effects observed in human data (Fig. 4f, right and Supplementary Fig. S3).

Our results show that the primacy and recency effects in sequential working memory can be generated from the cooperation of stable and flexible encodings of a neural circuit. In addition, stable and flexible encodings can be achieved simply from stable/flexible types of synaptic plasticity. Interestingly, stable and flexible encodings are two essential components among the working memory characteristics needed to retain and update information simultaneously. This implies that the serial-position effect reflects the most fundamental aspects of functional circuits for working memory.

### Working memory modulation by flexible/stable encoding balance

We observed that the coexistence of flexible and stable synapses generates the characteristic profile of sequential memory performance (Fig. 4). Furthermore, our model predicted that controlling the ratio of flexible/stable components would alter memory performance differentially by item order in the sequence (Fig. 5a); that is, stronger primacy effect would be observed when the ratio of stable synapse was increased (small *λ*), thus memory performance for early items would be improved (Fig. 5b). In contrast, weaker primacy effect would be generated when the ratio of flexible synapse was increased (large *λ*), thus memory performance would be worsened for early items. Therefore, memory performance could be altered item-order specifically by modulation of the flexible/stable synaptic balance. Specifically, average performance modulation by flexible/stable ratio control is predicted to be more significant for early-presented items (1^st^ ~ N-1^st^ items) than for the last item (N^th^ item) (Fig. 5c). This sequence-specific memory modulation effect is a key prediction of our sequential overwriting scenario with flexible and stable encodings.

**Figure 5.**
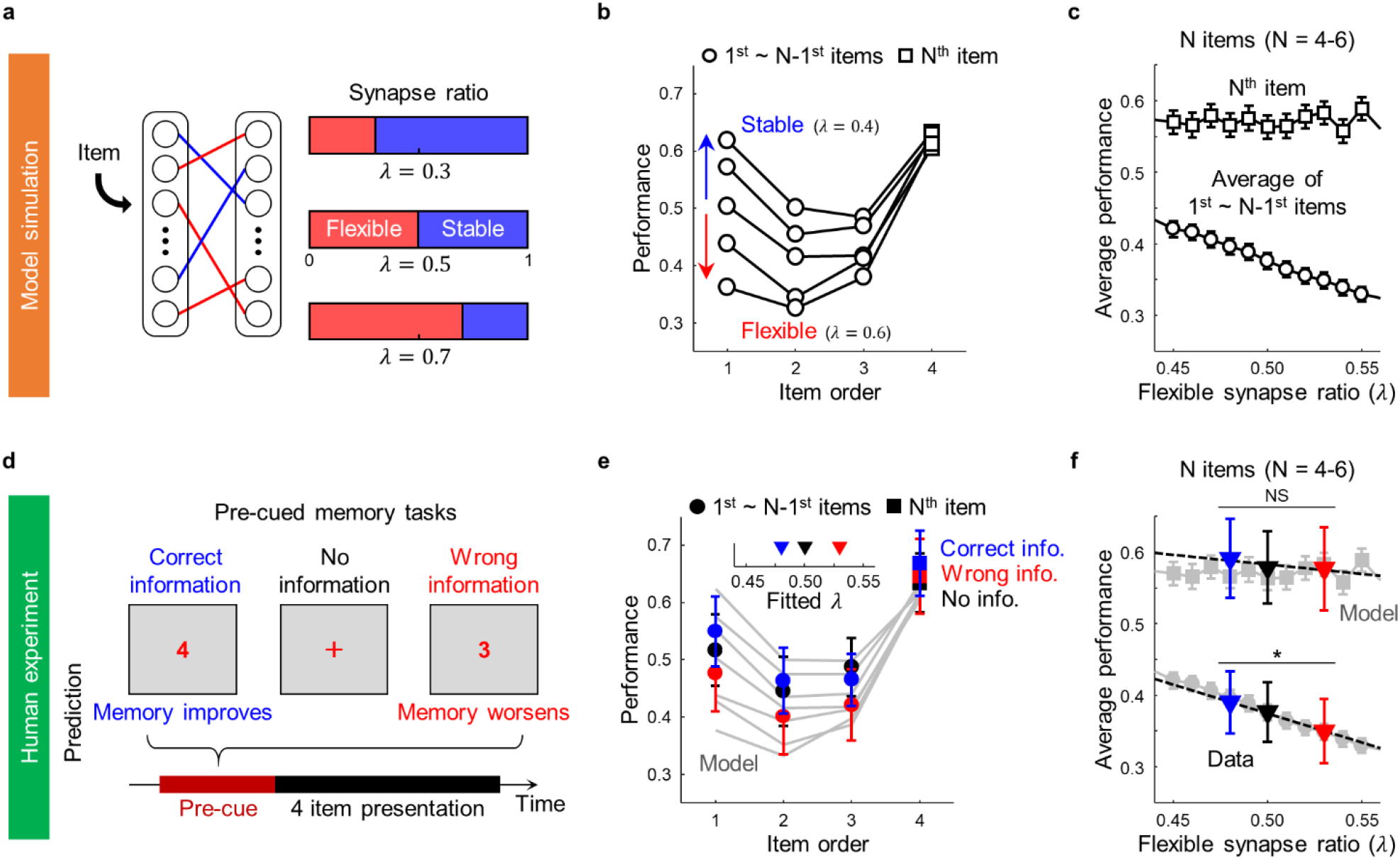
Change of flexible/stable synapse ratio modulates memory performance in sequence-specific way. **a**, Illustration of flexibility modulation in neural network model. **b**, Model simulation results of altered synaptic stability. Memory performance of early presented items (white circles) is improved if the network consists of more stable synapses (small *λ*), or worsened with more flexible ones (large *λ*). **c**, Memory performance changes with the flexible synapse ratio. The larger performance difference is observed in early presented items (white circles) rather than last items (white squares). **d**, Paradigm for pre-cued memory tasks. The three types of precues given: correct, no, and wrong information of the total number of items in a sequence. In the wrong information case, “N-1” is given before N items are presented. **e**, Sample result of a pre-cued memory task. Memory performance improves when the correct information is given (blue), while it worsens when the wrong information is given (red). Model performance with a different degree of flexibility (line) fit the observed performance (marker). (inset) Estimated flexibility from model fitting. **f**, Observed memory performance difference with estimated flexibility. As simulated in the model (gray markers), performance difference across conditions is better in early presented items than in the last item (for N-1 items, repeated measures ANOVA with Bonferroni post hoc correction, *F*(2,54) = 26.01, \**P* = 1.23 × 10 ^8^; for N^th^ item; *F*(2,54) = 1.30, *P* = 0.28). All error bars represent 95% confidence intervals.

With this hypothesis, we examined if this sequence-specific performance modulation could be observed in human data. Our hypothesis was that the flexible/stable encoding balance, could be altered if the degree of resource overwriting were changed, as shown in our model simulation (Fig. 2e and Fig. 3e). We might achieve this condition of overwriting variation by contrasting optimal/non-optimal allocation of memory resource. For this, we designed a human psychophysical experiment of memory allocation control. We hypothesized that memory allocation could be optimized if the total amount of information (or number of items) to memorize was given to the subjects prior to the test. For instance, if it were announced that “Four items will be given in the test”, then the subject could pre-estimate the size of resource allocation for each item that optimizes the degree of overlap between neural resources for different items. This pre-cue would effectively reduce the sequential overwriting in flexible coding and would increase the performance of early items in the sequence (Fig. 2e and Fig. 3e). On the contrary, if a wrong pre-cue were given so that the subject attempted non-optimal allocation of memory resource, the effect might be reversed and performance for early items degraded.

To test this idea, we performed a memory task with three pre-cue conditions: the total number of items was (1) correctly given (correct information), (2) not given (no information), or (3) wrongly given (wrong information), prior to item presentation (Fig. 5d). For wrong information, the number *N* – 1 was shown before *N* items were presented. Actually, *N* was varied from 4 to 6. As expected, memory performance was highest when the correct information was given and was lowest when the wrong information was given (Fig. 5e). In addition, performance difference was more noticeable in early items in the sequence. To estimate quantitatively the degree of flexible encoding from the experimental data, we simulated memory performance in the model network by varying the ratio of flexible synapses (*λ*) and by minimizing the mean squared error of performance between the model and data (Number of items = 4–6). As a result, the case of memory performance with correct information was well described by low flexibility conditions (*λ* = 0.48), while that of the wrong information case was well described by high flexibility conditions (*λ* = 0.53) (Fig. 5e and Supplementary Fig. S4). In addition, memory performance was largely altered in early presented items compared to the last items, as the model simulated (Fig. 5c and f). These results suggest that sequence-specific memory modulation by pre-cue could be described by manipulation of the flexible-stable encoding balance in the neural circuit.

## Discussion

In this work, we investigated a characteristic profile of the serial-position effect in sequential working memory, and proposed that the primacy and recency effects reflect stable and flexible encodings of neural circuits, both of which are required to retain and update information for working memory function. We also showed that flexible and stable memory function could be implemented by different types of synapses in a model neural network and that balance modulation of flexible/stable encoding could alter memory performance in a sequence-specific way.

Our new concept of sequential overwriting of memory resource provides a simple explanation for previous observations on working memory performance for simultaneous and sequential presentation of stimuli^42–44^, where memory performance is worse when stimulus information is presented sequentially than when it is presented simultaneously. This result has not been addressed by a simple resource model, because the total amount of allocated resources must be different across the presentation conditions, even though the number of stimuli was identical. Thus, the total amount of resource seems to vary for simultaneous and sequential presentations, which is controversial to the basic assumption of the resource model. Our sequential overwriting model, however, suggests that such a difference could arise from various conditions of resource overwriting. If there existed a resource overwriting, such that the resources for old items were degraded by a new item, the effect of overwriting would be noticeable only under the condition of sequential presentation, but not for simultaneous presentation. Thus, the observed difference between sequential and simultaneous presentation conditions was naturally understandable in our view. Another observation, that memory performance for the last item in sequential presentation is not different from that for simultaneous presentations^5^, also supports memory resource overwriting. In our model, the last item in a sequence is not affected by overwriting, and the performance must be the same as that in simultaneous presentation. Therefore, this experimental result is consistent with the prediction of our model (Fig. 2e).

From the fact that the last item in a sequence is not affected by resource overwriting, we were able to achieve another important finding directly from the observed human data: there is a profile indicative of memory resource allocation without resource overwriting (Fig. 3d). In this case, the amount of allocated resources decreases exponentially; thus we could investigate this primacy effect separately from the recency effect. Interestingly, this observation of declining resources is similar to the conceptual idea in previous models^18,25^, in which decreasing activation level or novelty-based encoding were suggested. The models assumed that early presented items are more strongly encoded than recent ones, consistent with the view of a stable coding scheme. Thus, our model suggests that the stable coding model provides a plausible mechanism for previous ideas of decreasing memory resources.

To provide an example of a neural circuit in which flexible and stable encodings contribute together, we simulated a model network with flexible and stable synapses (Fig. 4c and d). However, it is notable that collaboration of flexible and stable encodings can be achieved in numerous ways by other neural mechanisms, too. For instance, one study showed that intrinsic temporal stability of neuronal activity can be heterogeneous in a population, which may determine whether each neuron encodes information stably or dynamically^37^. In general, any neural parameters that modulate the stability of synaptic connection or activity could induce the combination of flexible and stable encodings at population level. It is also notable that distinct roles of flexible and stable coding have been observed in the memory function of flexible and stable values. A previous study reported that cells in the caudate nucleus encode values in two distinct forms^45^: neurons in the caudate head code flexible values while those in the caudate tail code stable values. Thus, the flexibility of encoding may vary by location, and probably by distinct types of neurons. Taken together, the collaboration of flexible and stable encodings could be generated by a variety of factors.

In summary, we found that the serial-position effect in sequential memory reflects distinct roles of flexible and stable encodings in neural memory circuits for working memory function. Our findings explain the origin of the serial-position effect and suggest that an association of flexible and stable encodings enables characteristic functions of working memory for retaining and updating information simultaneously. Our model provides a theoretical basis for understanding neural circuits for human working memory.

## Methods

### Subjects

Twenty-eight subjects (14 males, 14 females; 20 to 29 years old; all with normal or corrected normal vision) participated in the experiments after agreeing by written informed consent approved by the Institutional Review Board (IRB) of KAIST (KH2017-05). All procedures were carried out in accordance with approved guidelines.

### Sequential memory task and visual stimulus

Non-semantic visual patterns were used as a stimulus in the sequential memory task. Visual stimuli were blob-like patterns of within a 1.5° × 1.5° colored ring (bandwidth 0.1°, in visual space), and the pattern was generated as follows^46^:

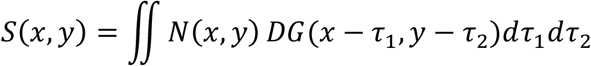

where *N*(*x,y*) is white noise and *DG*(*x,y*) is the difference between two 2D Gaussian filters (*σ*_1_ = 0.4°, *σ*_2_ = 0.8°, in visual space)(Supplementary Fig. S1a). A circular part of the pattern was normalized by z-scoring, and its absolute value was upper bounded by ‘3’. Subjects were positioned 160 cm away from the monitor and the visual patterns were presented on an LCD monitor (DELL U3014, 29.8 inch, resolution of 2560 × 1600, 60 Hz).

During the task, visual patterns (Number of items = 3-6) were presented sequentially at the center of the screen and subjects were asked to memorize their shape and order (Fig. 1a). A fixation cross at the center was presented in black (500 ms), red (1000 ms) and black (500 ms) in sequence, to inform the trial start. After a fixation screen (2000 ms), a stimulus was presented for 400 ms and an inter-stimulus-interval was given for 200 ms. After a 1000 ms delay, candidate patterns consisting of the presented stimuli and the same number of not-presented stimuli were given in a test session (Supplementary Fig. S1b). On the test screen, subjects were asked to recall freely the memorized sequence from the candidate patterns with a mouse click. They had to choose a sequential position and the item corresponding to that position.

Three conditions were tested in the pre-cued memory tasks (Fig 5d). Prior to the sequential presentation, the number of items to memorize was (1) correctly given, (2) not given, and (3) wrongly given to subjects to provide different conditions of pre-allocation of resources. In the wrong information case, a number that was one less than the actual number of presented items was given to subjects to induce miss-allocation of resource (e.g., ‘3’ was given as a pre-cue before ‘4’ items were presented). Twenty five percent of the pre-cue trials were under wrong information conditions. Three to six items were presented in the correct information and no information cases, and four to six items were presented in the wrong information case in random order. The ratio of trials for each condition was set as follows:

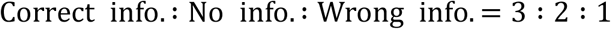

Subjects performed a training session (60 trials/session) and ten experimental sessions (72 trials/session). All codes for the experiment were generated with the MATLAB Psychtoolbox.

### Calculation of sequential memory performance

From the repetition of the sequential memory task sessions, the memory performance of each subject was measured. If the subjects chose both the item and order that matched the presented sequence, it was counted as correct. The performance in each order was then calculated. The response time of all trials was also measured, and its distribution for each condition was fitted with a log-normal function. The trials in which the response time lay outside of the 2σ (standard deviation) of the response time distribution were excluded from the analysis.

### Sequential overwriting model and memory resources

To model quantitatively the memory performance of the sequential memory task, we assumed that the amount of allocated resource *R_i_* determined the performance of the item positioned in order *i* as follows:

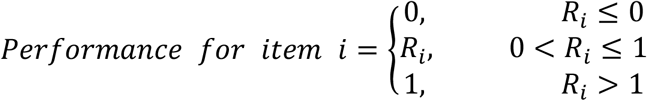

where *R_i_* is the amount of resource allocated for item *i*.

We also assumed that the previously allocated resources were overwritten when a new item was added as follows:

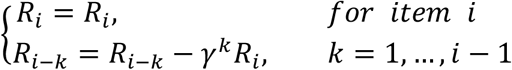

where *γ* is the overwriting ratio (Fig. 2) and *k* is the sequential distance between previous item and new item. Here, *R_i_* can be either identical (Fig. 2) or declining (Fig. 3) by order. To estimate the amount of allocated resources from the observed data (Fig. 3d), the performance for the last item was obtained from the average memory performance curve of the no-information condition, and fitted by

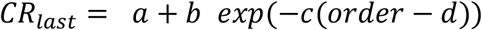

where *‘order’* varies from three to six.

### Data fitting with sequential overwriting model

To reproduce quantitatively the observed sequential memory performance, we fitted the model by minimizing the mean squared errors between the data and simulated curves. The amount of allocated resources at each order was estimated from the average memory performance of the last item. The fitted sequential overwriting ratio (*γ*\*) was searched from error minimization, using the “fmincon” function in MATLAB:

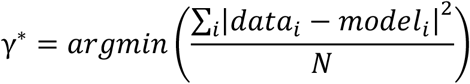

where *data_i_* (*models*) is the performance for order *i* in the data (model), respectively. *N* is the total number of items in a sequence (Fig. 3f).

### Neural network model

To study the neural basis of the serial-position effect, we used a model neural network that could learn and store sequential input patterns, adapted from a previous study^39^. The model network consisted of two layers: input and output layers (50 neurons each) of integrate-and-fire model neurons. Each input and output neuron was connected with a probability 0.2. In training sessions, six input patterns (10 Hz spike train for 100 ms) were given sequentially (50 times) and synaptic weights were updated using a STDP rule as follows:

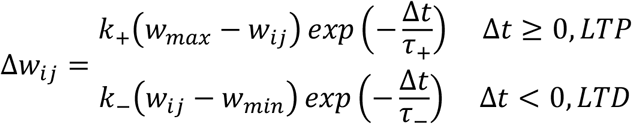

where Δ*t* = *t*_post_ – *t*_pre_ represents a spike timing interval. The other parameters set, were *k*+ = 0.6, *k*_−_ = −0.9, *τ*_+_ = 3 ms, and *τ*_−_ = 15 ms. Performance of the trained network (Memory index) was measured as the consistency of binary output spike patterns for the same inputs repeatedly given. A binary pattern was defined from the output firings: a number for each output neuron was set as ‘1’ if the neuron fired at least once during the repetitions, while it was set as ‘0’ if there was no response spike. Consistency was measured by averaging pairwise cross-correlations between all patterns of the repeated trials as follows (see previous Methods^39^ for details):

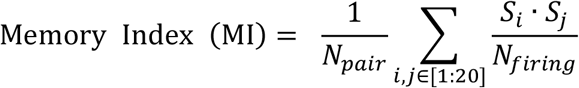

where *S_i_* represents the *i*^th^ binary pattern of output firing, *N_pair_* denotes the number of all pairs, and *N_firing_* is the total number of fired output neurons. To rescale a memory index into memory performance, we applied a sigmoid function to memory index as a response transfer function (Supplementary Figure S2c), based on the observation that behavior results could be described by a logistic function ogf neural activity^47^. We used the sigmoid function as follows:

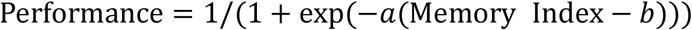

Each constant was estimated by minimizing the mean squared error of the sequential memory performance between data and model using the MATLAB function ‘fmincon’ (*a* = 6.78, *b* = 0.34 for flexible synapse only case; *a* = 14.92, *b* = 0.59 for stable synapse only case; *a* = 17.5, *b* = 0.46 for both flexible and stable synapse case). For learning, we defined two types of synapses: flexible and stable synapses. For the flexible one, synaptic weight was allowed to change continuously during the entire learning event. In contrast, for a stable one, the synaptic weight was set to remain unchanged when the weight saturated to 99% of the maximum or minimum value.

### Statistical test

The type of statistical test and corresponding p-values used in the analysis were given in figure captions and the main text. One-way ANOVA with Bonferroni correction was used to examine performance differences across the pre-cue conditions.

## Supplementary information

Supplementary figures and legends are available in Supplementary Information.

## Acknowledgements

This work was supported by the National Research Foundation of Korea (NRF) grant funded by the Korea government (MSIT) (No. NRF-2016R1C1B2016039, NRF-2017R1E1A2A02080940) (to S.P.).

## Author contributions

H.L. designed and performed the psychophysics experiments, analyzed data, and wrote the manuscript. W.C. and Y. P. analyzed data. S.P. conceived the project, directed the experiments and wrote the manuscript. All authors discussed, commented on and revised the manuscript.

## Competing interest declaration

Authors declare no competing interests.

---

**Supplementary Figure S1.**
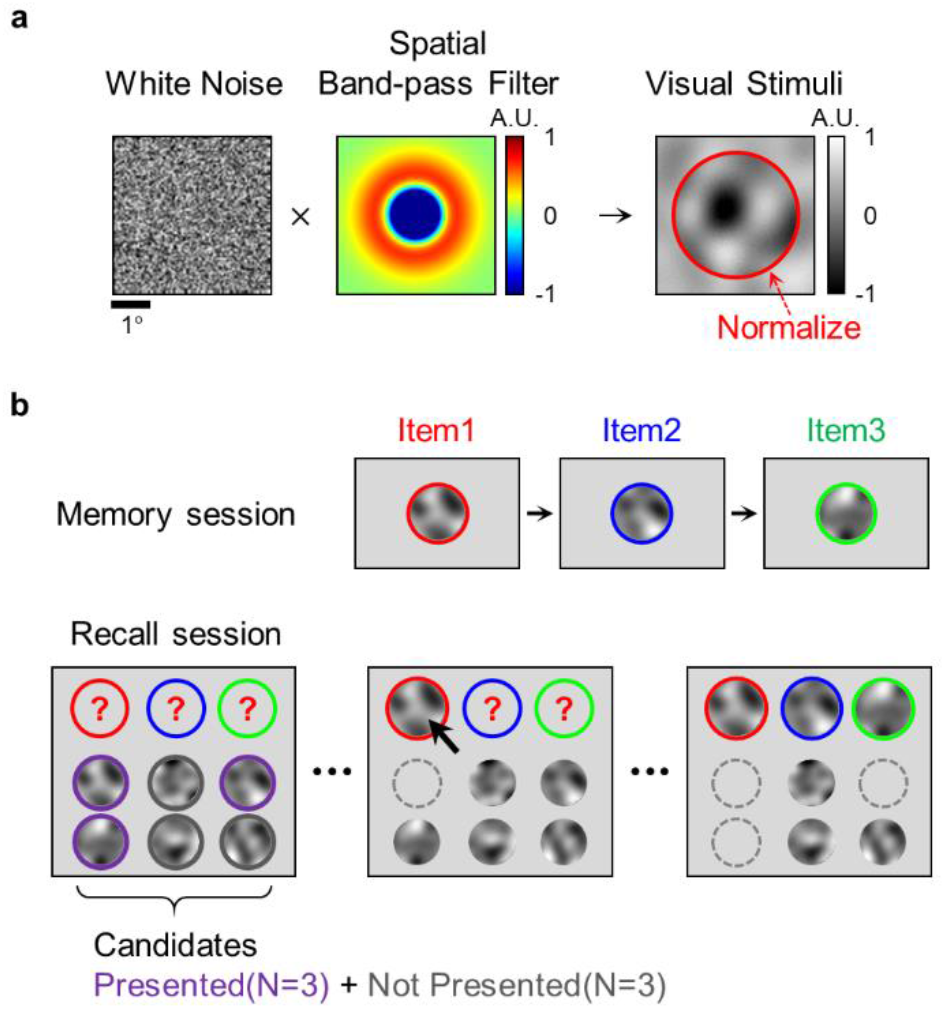
Stimulus generation and paradigm of sequential memory task. **a**, Design of visual stimuli. Visual stimuli are generated by filtering a two-dimensional white noise with a spatial band-pass filter. The filter is designed with a difference between two Gaussian distributions (σ_1_ = 0.4°, σ_2_ = 0.8°, in visual space; see Methods for details). **b**, Design of sequential memory task. Subjects memorize sequentially presented items (N_items_ = 3-6) during memory session. In recall session, subjects choose the presented items and their presented order among candidates. Items presented during the memory session (purple circle) and the same number of not-presented ones (gray circle) were given as candidates.

**Supplementary Figure S2.**
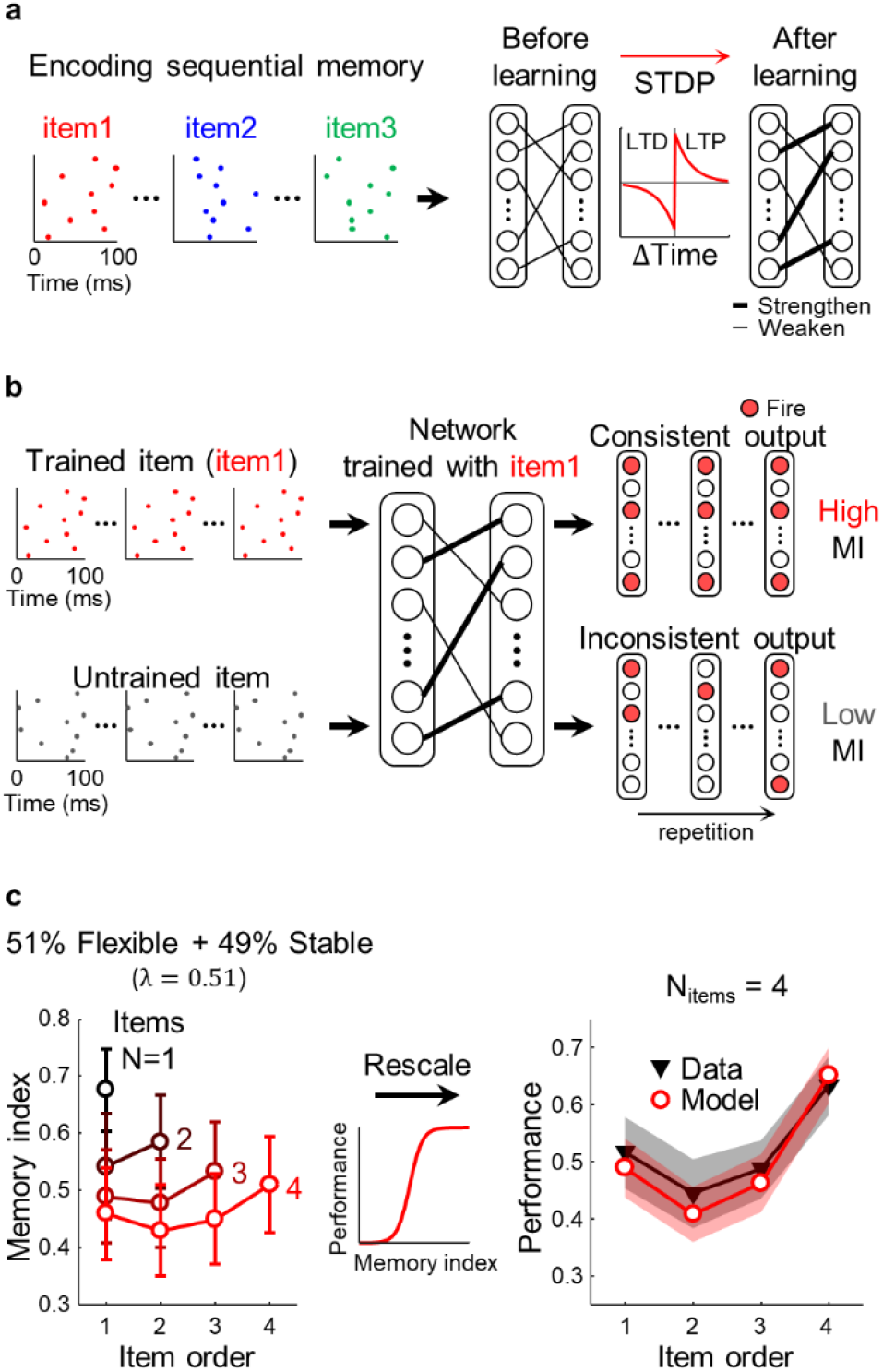
Paradigm of neural network simulation. **a**, Scheme of memory encoding. Sequential spike trains are encoded in a neural network using an STDP learning rule. Each spike train (item) consists of a spike at random timing within 100 ms and is fed into a network 50 times, for 5 s. **b**, Scheme of memory test. The consistency of output firing patterns is measured for the repeatedly given items and defined as a memory index (see Methods for details). The memory index is high for the trained items (top), while it is low for the untrained items. **c**, Simulated results of the model (bottom). (left) Memory index by item order. Four memory index curves show how the network responds to trained items, as the number of items in a sequence increase (from N_items_ = 1-4). The U-shaped memory performance curves are observed after four items are encoded. (middle) To rescale the memory index into memory performance, a sigmoid function is applied. (right) Comparison of performance between the experimental data and model (N_items_ = 4). Our model generates the serial-position effect observed in the experiment. Shaded area represents 95% confidence intervals.

**Supplementary Figure S3.**
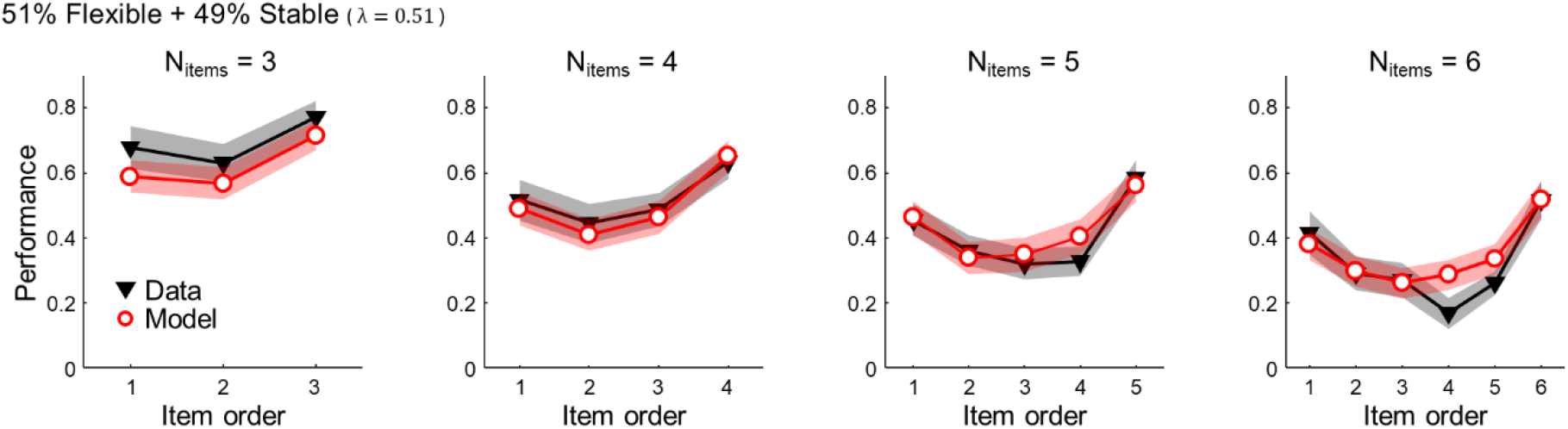
Results of model fitting for different numbers of items. The neural network regenerates the serial-position effects observed in the human psychophysical experiments. The red line represents the simulated results of the model, while the black line represents the experimental data. Shaded area represents 95% confidence intervals.

**Supplementary Figure S4.**
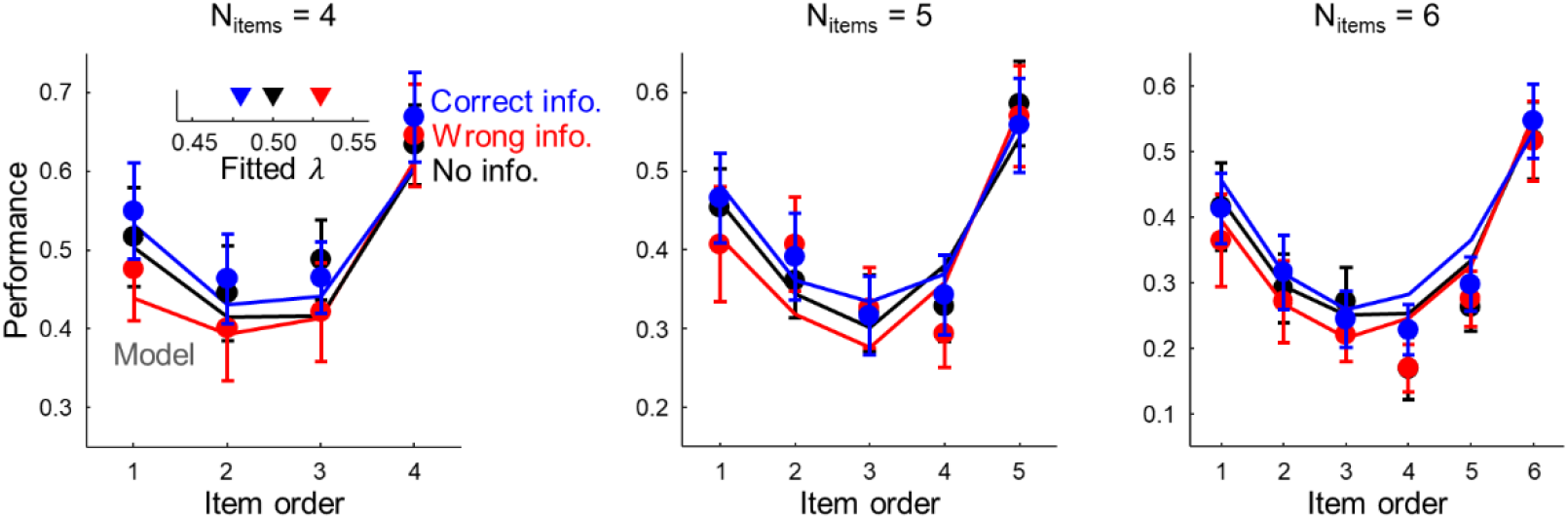
Results of pre-cued memory task and model fitting. A neural network with low flexible synapse ratio is able to generate the memory performance of the correct information case (blue), while that with a high flexible synapse ratio is able to generate the performance of the wrong information case (red). Error bars represent 95% confidence intervals.

